# Effective repair of joint cartilage using human pluripotent stem cell-derived tissue

**DOI:** 10.1101/340935

**Authors:** Oliver F. W. Gardner, Subhash C. Juneja, Heather Whetstone, Yulia Nartiss, Jakob T. Sieker, Christian Veillette, Gordon M. Keller, April M. Craft

**Author notes:** These authors contributed equally to this work (Oliver F. W. Gardner, Subhash C. Juneja). Corresponding author: April M. Craft, Ph.D., Dept. of Orthopaedic Surgery & Research, Mailstop 3096, Enders Research Building, Suite 260, Boston Children’s Hospital, Boston, MA 02115.

## Abstract

Adult articular cartilage lacks significant regenerative capacity, and damage to this tissue often leads to progressive joint degeneration (osteoarthritis). We developed strategies to generate articular cartilage from human pluripotent stem cells (hPSCs) as a source of clinically relevant tissues for joint repair^1^. Previously, we demonstrated that these chondrocytes retain cartilage forming potential following subcutaneous implantation in mice. In this report, we evaluated the potential of human embryonic stem cell (hESC)-derived articular cartilage tissue to repair osteochondral defects created in the rat knee. Following implantation, the hESC-derived cartilage maintained a proteoglycan and type II collagen-rich matrix, and was well integrated with native rat tissue at the basal and lateral surfaces. The ability to generate cartilage tissue with integrative and reparative properties from an unlimited and robust cell source represents a significant clinical advance for cartilage repair that can be applied to large animal models and ultimately to patient care.

---

Using serum-free directed differentiation methods previously described ^1^, we generated articular cartilage-like tissue containing abundant proteoglycans (Safranin-O positive) and type II collagen but lacking type X collagen from the HES2-RFP hESC line which constitutively expresses red fluorescent protein (RFP, Figure 1A-D)^2^. We generated two osteochondral defects (1.45 mm diameter) in the trochlear groove of the rat femur, press-fitted size-matched biopsy punches of hESC-derived tissue into the defects, and sealed them with fibrin glue (Figure 1E-F). Six to twelve weeks later, we assessed graft retention and cartilage quality compared to control defects treated with fibrin glue alone (Figure 1G-H).

**Figure 1.**
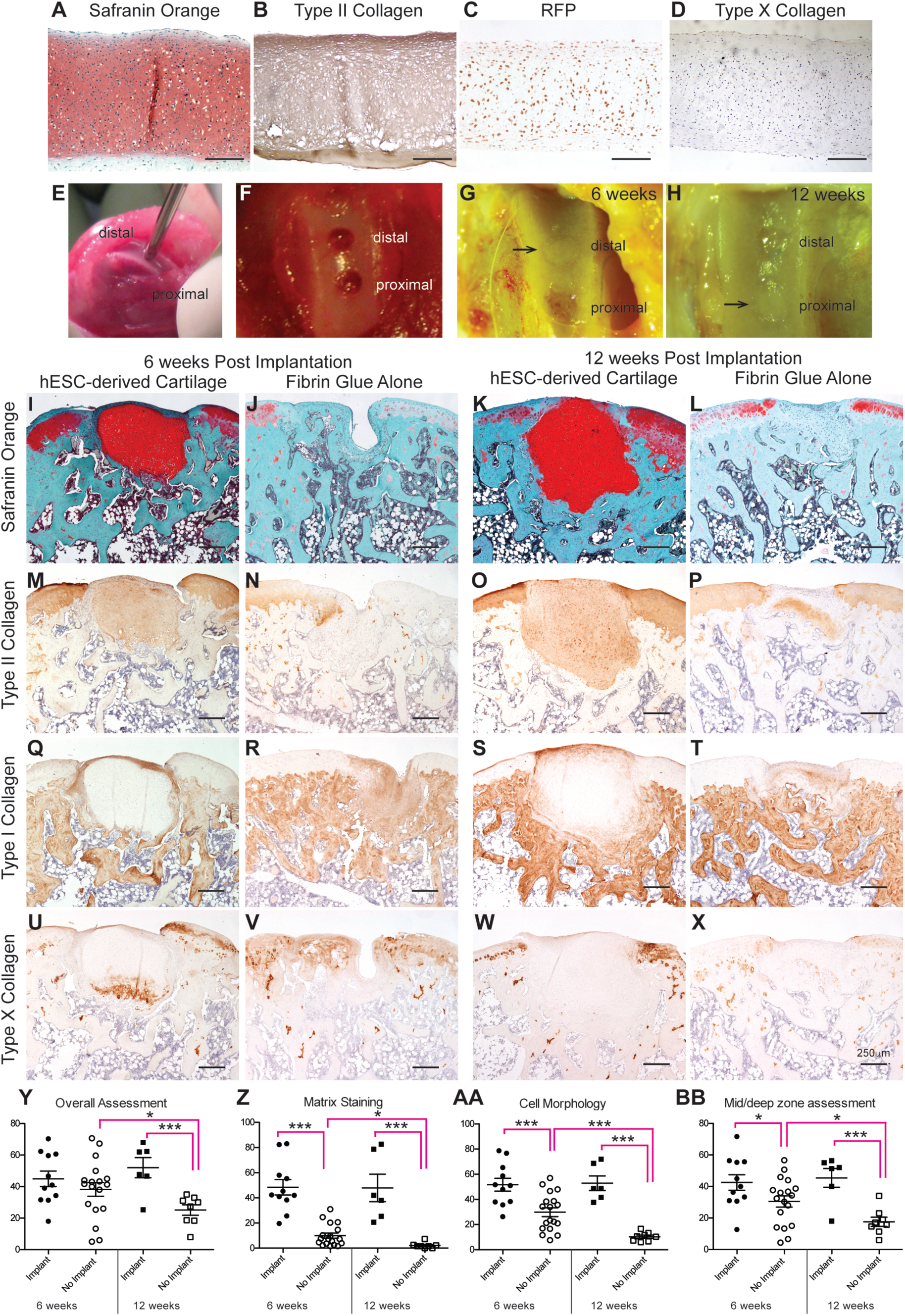
Improved cartilage repair following implantation of hESC-derived articular cartilage tissues into osteochondral defects created in the rat knee. (A-D) Pre-implantation cartilaginous tissue derived from the HES2-RFP hESC line stained positive for Safranin O (indicating proteoglycans), type II collagen and RFP, but not type X collagen. (E-F) Two 1.45 mm defects were created ~0.5 mm apart in the femoral trochlea groove of the right knees of athymic nude rats. Macroscopic images of defects that were implanted with hESC-derived tissue after 6 (G) or 12 weeks (H), arrows indicate defects in which human tissue was identified histologically. (I-X) Histological and immunohistochemical staining of representative implanted and control defects at 6 and 12 weeks post-implantation. Scale bar represents 200 µm (A-D), 250 µm (I-X). (Y-BB) Outcome assessments based on histological scoring according to a modified ICRS II system (see methods for description). Scores for each randomized defect were averaged from three blinded reviewers and plotted vertically according to treatment (implanted defects are represented by filled icons, outlined icons represent control defects) and time (defects after 6 weeks are represented as circles and defects after 12 weeks are squares). The mean and standard deviation are indicated. Scores for remaining ICRS II parameters are shown in Supplementary Figures 1 and 2. *p<0.05, **p<0.01, ***p<0.001.

Eleven of the twenty-two defects analyzed after 6 weeks, and six of the eighteen defects analyzed at 12 weeks contained human cells/tissue. Absence of human cells in some of the defects was likely due to delamination of the tissue upon implantation, an inherent risk of this type of study ^3^. Defects containing hESC-derived tissues contained significant regions of proteoglycan- and type II collagen-rich cartilage tissue after 6 and 12 weeks (Figure 1I-P). In contrast, control defects generally lacked proteoglycans and type II collagen, but were enriched for type I collagen, indicating the presence of fibrocartilage-like tissue (Figure 1R, T). Type I collagen was also present in host tissue at the periphery of the human tissue implant and at the articular surface (Figure 1Q, S). Type X collagen, a marker of chondrocyte hypertrophy, was present in 5 of the 11 defects at 6 weeks and 1 of 6 defects at twelve weeks (representative defects shown in Figure 1U and 2F), with the highest levels localizing to the subchondral interface between implant and host tissue.

**Figure 2.**
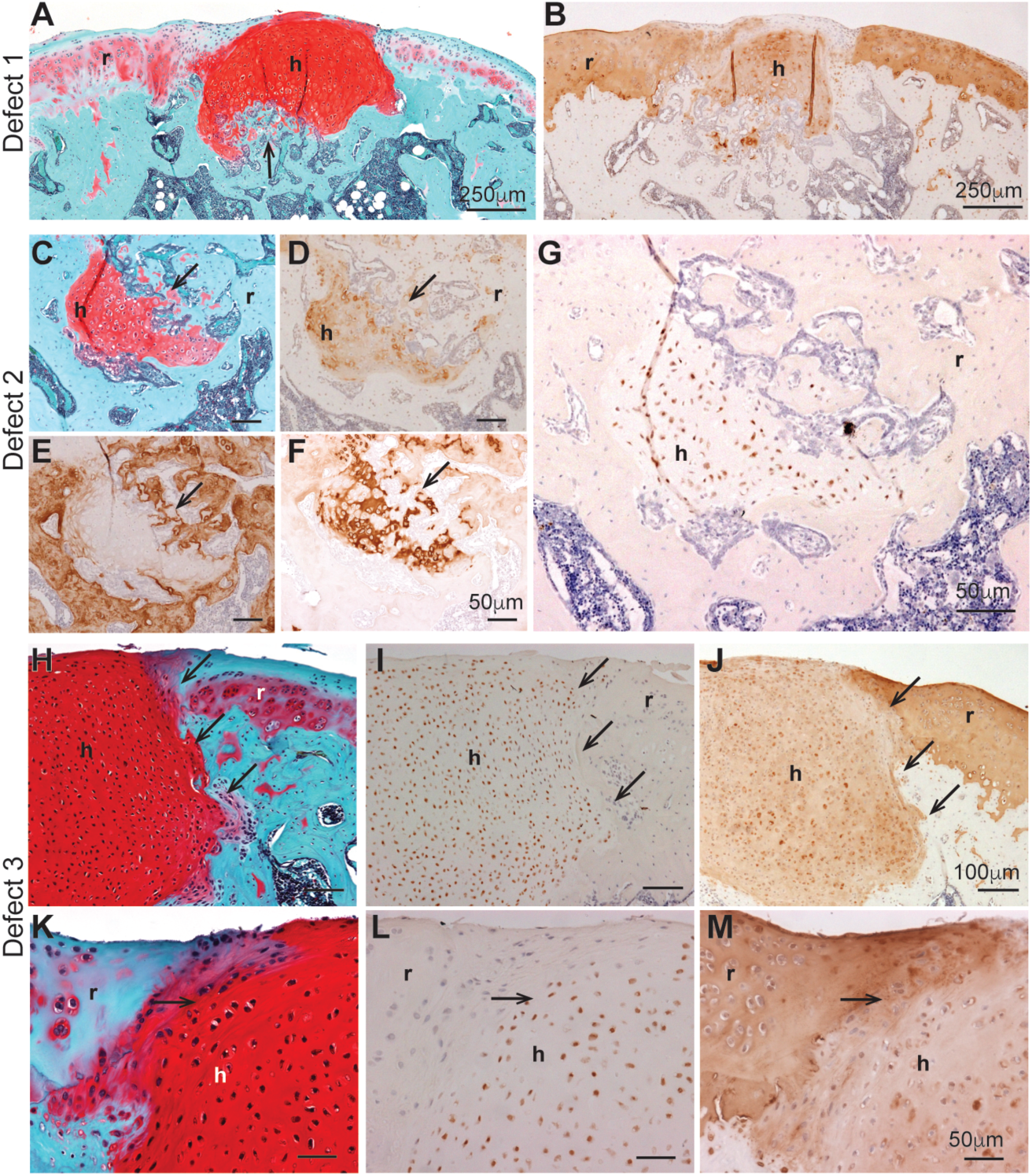
Evidence of integration and remodeling of hESC-derived cartilage implants in the rat knee. Three representative defects are shown from two different animals at 6 weeks post-implantation (A-G) and one animal at 12 weeks post-implantation (H-M). Defect 1 (A, B) shows cartilage implant congruent with the native rat cartilage at the articular surface. Defect 2 shows cartilage implant encased within subchondral bone (C-G). An arrow indicates likely orientation of remodeling (A, C). hESC-derived cartilage implants stain positively with Safranin O (A, C, H, K) and type II collagen (B, D, J, M). Type I collagen/bone matrix (E) was found directly adjacent to the human tissue in Defect 2. Human cells (RFP-positive, G) were identified only within the calcified cartilage regions marked by type II (D) and type X (F) collagen, and were not found in the adjacent bone tissue (E). Representative Defect 3 at 12 weeks post implantation shows basal and lateral integration of hESC-derived cartilage tissue implant in the absence of bone formation (H-M). The Safranin O-positive hESC-derived cartilage implant was congruent with the surrounding native rat cartilage (H, K). Human cells, identified by the expression of RFP, were laterally integrated within the host rat tissue (I, L, arrows). Note the uninterrupted type II collagen network between the hESC-derived cartilage implant and the native rat cartilage (J, M). Scale bar represents 250 µm (A, B), 50 µm (C-G; K-M), 100 µm (H-J). Abbreviations: r, rat; h, human.

We quantified cartilage repair in defects receiving implants versus controls using a modified International Cartilage Repair Society (ICRS) II scoring system^4, 5^ where 0 represents poor repair and 100 represents normal cartilage. The overall assessment scoring parameter was not significantly different between the defects receiving implants and controls after 6 weeks, but defects with human tissue scored significantly higher at 12 weeks (Figure 1Y). Implanted defects also had significantly higher scores in matrix staining, cell morphology, and middle/deep zone assessment at both 6 and 12 weeks. Scores for tidemark formation and vascularization significantly increased from 6 weeks to 12 week in defects receiving implants, while scores for matrix staining, cell morphology, middle/deep zone, and overall assessment significantly decreased in control defects over time (Figure 1Z-BB, Supplementary Figures 1 and 2). These results indicate that cartilage repair quality was superior in defects receiving hESC-derived implants compared to those receiving fibrin glue alone.

Finally, we evaluated the responses of the implanted human tissue and the host rat tissue via histology. Implanted tissue was well integrated basally and laterally with host tissue at both time points (Figure 2). Remodeling of human tissue typically occurred in areas that were exposed to the marrow cavity, similar to the process of endochondral bone formation^6^. Cells in these regions increased in size, alignment (Figure 2A, C, arrows) and type X collagen expression (Figure 2F), while proteoglycans and type II collagen abundance persisted (Figure 2A-D). Some defects with advanced remodeling contained small regions of unusually high levels of proteoglycans and type II collagen surrounded by type I and X collagen (Figure 2C-F, arrows), a phenomenon indicative of *de novo* bone formation^7^. RFP positive cells persisted in the proteoglycan-rich areas but not in the surrounding bone (Figure 2G), indicating that the implanted human tissue was being remodeled, providing a scaffold for the development of host bone. Remodeling of the implant appeared to facilitate integration (e.g., Figure 1I, 2A), and likely protected against graft delamination. At the articular surface and in regions where human tissue did not come into contact with bone marrow, we observed a continuity of matrix across clear boundaries between host cells and implanted cells (Figure 2H-M).

In this report we have demonstrated the ability of the hESC-derived articular cartilage tissues to engraft, persist and functionally integrate in an orthotopic site. The results of this study demonstrate that this is a viable approach for repairing damaged cartilage. Future studies in large animal models will explore the potential to treat osteochondral defects with a single hESC-derived cartilage implant.

## Supplemental Material

Supplemental information including methods follows this manuscript.

## Acknowledgements

The authors would like to thank Benedikt Proffen and Johannes Konrad (Boston Children’s Hospital) for histological scoring and members of the Craft laboratory for critical reading of the manuscript. This work was supported by an Accelerating Discoveries Research Award from the McEwen Centre for Regenerative Medicine, the Arthritis Program at the Krembil Research Institute, and a generous gift from the Krembil Foundation.

---

**Supplementary Figure 1.**
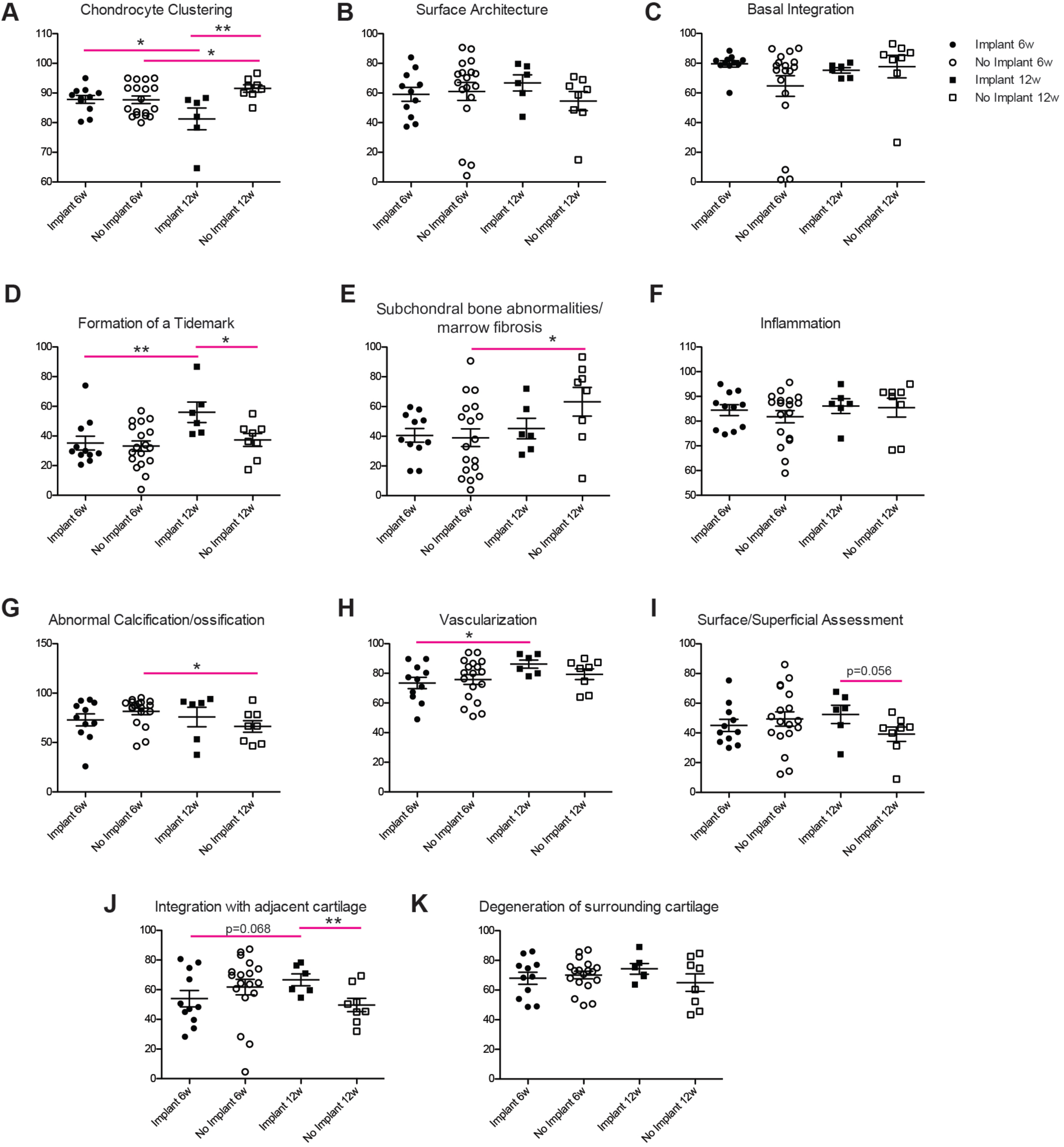
Outcome assessments of all other parameters comparing implanted defects for which human cells were detected and control defects which were treated with fibrin glue alone. *p<0.05, **p<0.01, ***p<0.001.

**Supplementary Figure 2.**
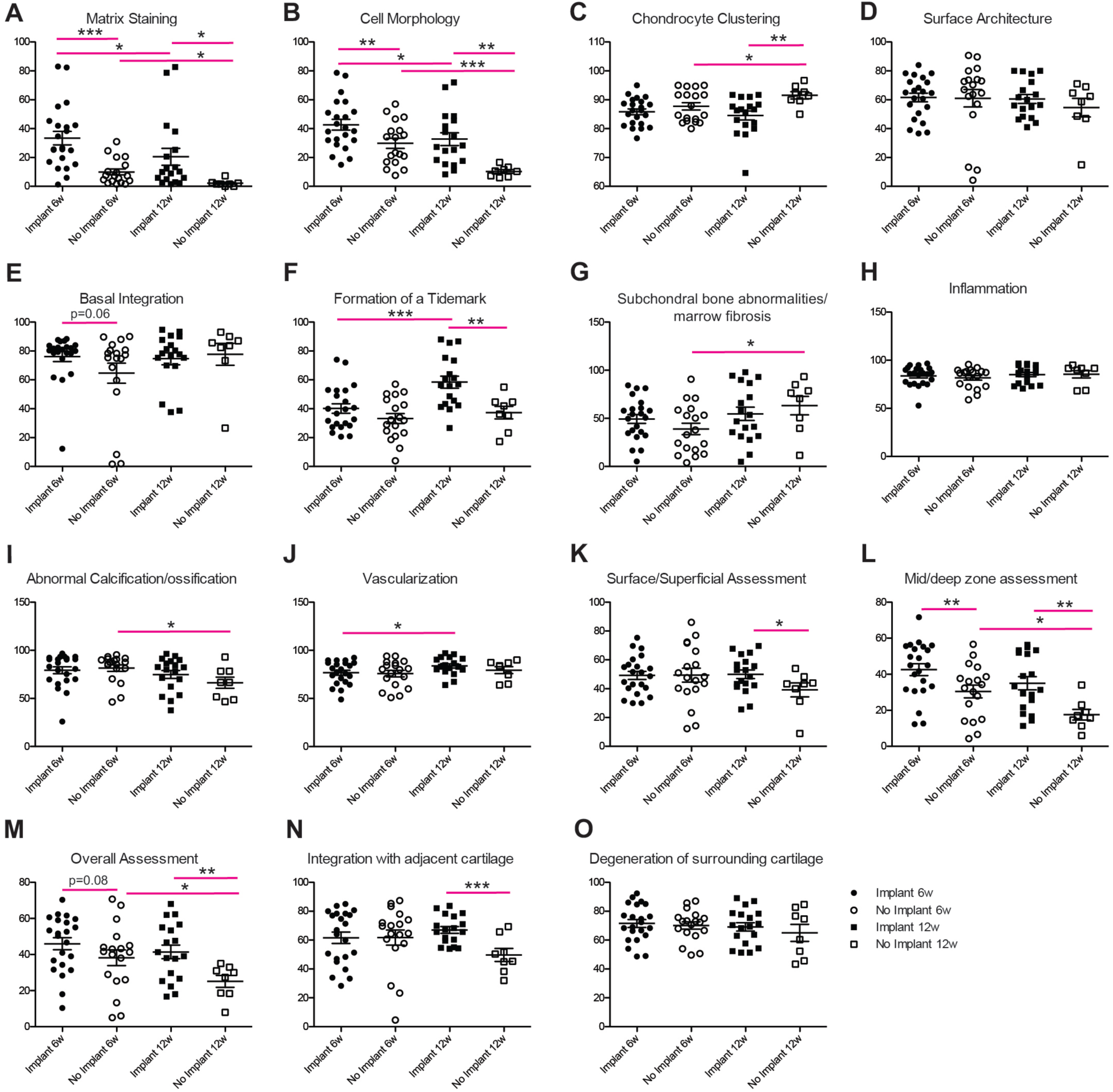
Outcome assessments of all parameters and all defects, including those in which human tissue was implanted but human cells were not identified in subsequent histological analysis. *p<0.05, **p<0.01, ***p<0.001.

## Materials and Methods

### Cell culture and tissue preparation

Articular cartilage tissues were generated in serum-free conditions as described previously ^1^ from the HES2-RFP hESC line that constitutively expresses the red fluorescent protein (RFP) ^2^. Briefly, hESCs (mycoplasma-free) were cultured as embryoid bodies for 3 days in the presence of Activin A, BMP4 and bFGF (R&D Systems) to induce early stage mesoderm formation. Chondrogenic mesoderm was induced in monolayer with an inhibitor of type I activin receptor-like kinase (ALK) receptors SB431542 (Sigma) (5.4 µM), bFGF (20 ng/ml) and the type I BMPR inhibitor Dorsomorphin (Sigma) (4 µM) until day 15. Cartilage tissues were generated in micromass format and cultured for an average of 14 weeks in chondrogenic media: high glucose DMEM (Gibco) supplemented with 1× ITS supplement (Gibco), ascorbic acid (Sigma) (50 µg/ml), proline (Sigma) (40 µg/ml) and dexamethasone (Sigma) (0.1 µM) and TGFβ3 (R&D Systems) (10 ng/ml).

### Surgical procedure

All animal studies were approved by the Animal Resources Committee at University Health Network, Toronto, Ontario, Canada. Homozygous athymic nude rats (NIH nude rats, Crl:NIH-Foxn1^rnu^; *rnu/rnu*) were purchased from Charles River Laboratories and were provided with sterile housing. Surgeries were conducted on animals at eight to nine weeks of age. Animals were anesthetized under 5% isoflurane and maintained at 2% isofluorane during surgery. Buprenorphine (0.03 mg/kg) was used as an analgesic drug. Pre-operative and post-operative care was provided to animals according to the protocol.

The right knee area was shaved and cleaned with iodine surgical scrub (7.5% iodine), 70% isopropanol and 10% Providone iodine (equivalent to 1% iodine) in sequence. A longitudinal incision was made on the anterior aspect of knee with a surgical scalpel blade no.15 to expose the knee joint. The skin at medial and lateral side of knee joint was dislodged using two backward strokes with a pair of surgical scissors. A second cut was made on the knee joint medially on the muscle layer. The muscle layer along with the quadriceps tendon was pushed laterally to expose trochlear region of femur.

Two osteochondral defects (0.5 mm apart, one proximal, one distal) were created in the femoral trochlear groove with the help of a sorer (1.45 mm diameter, Fine Scientific Tools, Inc.). Animals were assigned at random to the experimental (implant) or control group. Size matched biopsy punches of hESC derived cartilage tissue were press-fitted into defects with the help of cotton gauze, and then sealed with fibrin glue (TISSEEL, Baxter). Control defects were filled with fibrin glue alone. A total of 26 control defects were created in 13 knees, hESC-derived cartilage tissue was implanted into 40 defects in 20 knees.

The muscle layer was reverted back to the medial side and quadriceps tendon was placed in its original position. The medial muscle layer was sutured discontinuously at three points with 5-0 Polysorb™ suture (Covidien^*^). The skin was closed using intradermal continuous suturing using MONOCRYL^*^ suture (Ethicon), and glued with a thin layer of tissue adhesive (Vetbond, 3M). Following surgery, animals were monitored according to approved animal protocol, and then permitted to move freely for 6 or 12 weeks.

### Tissue Retrieval and Histology

Immediately following euthanasia, the knee joints were excised, and the defects were exposed and imaged macroscopically. Whole joints were then fixed in 10% neutral buffered formalin (Sigma) and decalcified for approximately ten days in Immunocal Decalcifier solution (DECAL Company) with changes every three days before being processed for paraffin embedding and histological analyses. Sagittal sections (5 µm thickness) of the trochlear groove of the femur containing defect(s) were stained with 0.1% Safranin O (Electron Microscopy Sciences) and counterstained with Weigert’s haematoxylin (Electron Microscopy Sciences) and 0.02% fast green (Electron Microscopy Sciences). Immunohistochemistry was used to visualize type I (Sigma), II (clone 6B3 EMD Millipore) and X collagen (Clone X53, Quartett, Germany) and red fluorescent protein (ab34771, Abcam). The M.o.M™ (Mouse on Mouse) Detection Kit (Vector Laboratories Inc.) was used with the type I and type II collagen monoclonal antibodies to minimize background staining. Detection was performed using VECTASTAIN ABC and DAB reagents (Vector Laboratories Inc.). Sections were counterstained with Meyers Haematoxylin (Electron Microcroscopy Sciences). Images were acquired using a Nikon Eclipse 80i microscope and a Nikon Digital Sight DS-Ri1 camera.

### Histological Scoring & Statistical Analysis

Quality of cartilage repair was determined using a modified ICRS II scoring system ^3^. The ICRS II scoring system was modified in the following ways: Tissue Morphology parameter was removed due to the requirement to view under polarized light, and this requirement was also removed from Middle/Deep zone assessment parameter. In addition, the ‘integration with adjacent cartilage’ and ‘degeneration in native adjacent cartilage’ parameters were added from the O’Driscoll scoring system ^4^.

Representative Safranin O-stained sections of each defect were randomized and scored by three independent, blinded individuals. Each defect was scored for matrix staining, cell morphology, chondrocyte clustering, surface architecture, basal integration, formation of a tidemark, subchondral bone abnormalities/marrow fibrosis, inflammation, abnormal calcification/ossification, vascularization, surface/superficial assessment, mid/deep zone assessment, overall assessment, integration with adjacent native cartilage, and degeneration of adjacent native cartilage. Each parameter was scored from 0 (poor repair) to 100 (normal articular cartilage).

For each of the fifteen microscopic parameters assessed, the scores from three blinded reviewers were averaged for each defect and plotted per treatment and time point. In order to compare successfully implanted defects to unimplanted defects, we excluded from these analyses the defects that received hESC-derived tissue but for which human tissue was not detected by immunohistochemical staining. A summary of outcome assessments of all defects, including those in which human tissue was implanted but not detected histologically, is provided as Supplementary Figure 2.

Statistical analyses were performed in Prism (GraphPad) using type 2 one-tailed t-tests to compare defects receiving implants to those that did not receive implants at each time point, and to compare implanted or unimplanted defects over time. A p-value of less than 0.05 was considered significant.

